# Targeting long non-coding RNA *NUDT6* enhances smooth muscle cell survival and limits vascular disease progression

**DOI:** 10.1101/2022.11.14.516372

**Authors:** Hanna Winter, Greg Winski, Albert Busch, Ekaterina Chernogubova, Francesca Fasolo, Zhiyuan Wu, Alexandra Bäcklund, Bohdan B. Khomtchouk, Derek J. Van Booven, Nadja Sachs, Hans-Henning Eckstein, Ilka Wittig, Reinier A. Boon, Hong Jin, Lars Maegdefessel

## Abstract

Long non-coding RNAs (lncRNAs) orchestrate various biological processes and regulate the development of cardiovascular diseases. Their potential therapeutic benefit to tackle disease progression has recently been extensively explored. Our study investigates the role of lncRNA Nudix Hydrolase 6 (*NUDT6)* and its antisense target Fibroblast Growth Factor 2 (*FGF2)* in two vascular pathologies: abdominal aortic aneurysms (AAA) and carotid artery disease. Using tissue samples from both diseases, we detected a substantial increase of *NUDT6*, whereas *FGF2* was downregulated. Targeting *Nudt6 in vivo* with antisense oligonucleotides in three murine and one porcine animal models of carotid artery disease and AAA limited disease progression. Restoration of FGF2 upon *Nudt6* knockdown improved vessel wall morphology and fibrous cap stability. Overexpression of *NUDT6 in vitro* impaired smooth muscle cell (SMC) migration, while limiting their proliferation and augmenting apoptosis. By employing RNA pulldown followed by mass spectrometry as well as RNA immunoprecipitation, we identified Cysteine and Glycine Rich Protein 1 (CSRP1) as another direct *NUDT6* interaction partner, regulating cell motility and SMC differentiation. Overall, the present study identifies *NUDT6* as a well-conserved antisense transcript of *FGF2. NUDT6* silencing triggers SMC survival and migration and could serve as a novel RNA-based therapeutic strategy in vascular diseases.

## Introduction

Atherosclerosis, the underlying pathology of most cardiovascular diseases, remains the most prominent cause of death worldwide (1). Advanced atherosclerosis in carotid artery plaques is the most common reason for ischemic forms of large artery stroke while progressing abdominal aortic aneurysms (AAAs) are closely associated with the atherosclerotic burden and vascular inflammation within the infrarenal section of the aorta. Both diseases share similar molecular characteristics, including endothelial dysfunction (2,3), enhanced monocyte infiltration and macrophage activity (4,5), increased smooth muscle cell (SMC) apoptosis (6,7), as well as impaired extracellular matrix (ECM) remodeling (8,9). In AAAs, these mechanisms can culminate in an acute rupture with mortality rates above 80% (10,11). Vulnerable atherosclerotic lesions in the carotid artery can become unstable and rupture. This in consequence, leads to transient ischemic attacks (TIAs), *amaurosis fugax*, and ultimately ischemic stroke, causing physical impairments and death (12).

Apart from sharing similar pathophysiological features, both diseases also have similar risk factors, including dyslipidemia, arterial hypertension, family history/genetic association, advanced age, male sex, and smoking (13). Of further importance, about 60% of all AAA patients die from other cardiovascular diseases, such as stroke or myocardial infarction, again emphasizing the overlap in risk factor profiles and potentially shared underlying mechanistic drivers of both diseases (8,14). While several medications were discovered to slow down the atherosclerotic process in the carotid artery limiting the risk of plaque rupture, no treatment (apart from surgical intervention) is currently available for AAA patients to slow down expansion and reduce their risk of an acute rupture (15). Clinical handling of AAA patients is further hampered by the fact that currently, no tools (*e*.*g*., biomarkers) exist that allow foreseeing growth rates and rupture risk for any given individual.

SMC dynamics and fate in both pathologies play crucial roles during disease exacerbation. Under physiological conditions, vascular SMCs regulate the tone and contraction of arteries, thus maintaining the morphological integrity of the vasculature (16). When SMC survival is impaired, the vessel wall and plaque fibrous cap weaken, leading to lesion instability and aortic dilatation (6,9). When SMCs become hypo-migratory, fibrous cap thickness is decreased, and the aneurysm diameter expands (6,17). Furthermore, augmented matrix production and states of limited proliferation in SMCs accelerate the progression of both diseases (18,19).

Natural antisense transcripts (NATs) are a subclass of long non-coding RNAs representing the largest group of RNAs in mammalian genomes (20). NATs are generally transcribed from the opposing strand to protein-coding genes. For most NATs, a repressive function on the expression of their partner sense transcript has been described in *cis* (21). This becomes interesting in diseases where the upregulation of classically considered ‘undruggable’ targets, such as transcription or growth factors, is desirable (22,23). However, targeting the NAT that blocks transcription increases their expression levels, thus imposing potential benefits as future therapeutic approaches. Further, NATs were shown to exert their action faster than transcription factors due to their spatial proximity which allows for immediate adaptation (24,25).

In this current study, we have discovered that the lncRNA Nucleoside diphosphate-linked moiety X motif 6 (*NUDT6*), a NAT to the fibroblast growth factor 2 (*FGF2*), negatively affects carotid artery plaque stability as well as AAA progression. *NUDT6* overexpression limits survival, proliferation, and migration in arterial SMCs. Conversely, its silencing by using site-specific antisense oligonucleotides (GapmeRs), limits plaque rupture rates and experimental AAA growth in four preclinical disease models, thus providing an avenue for developing new therapeutic strategies to limit the burden of both vascular pathologies.

## Results

### *NUDT6* and *FGF2* are deregulated in fibrous caps of advanced human atherosclerotic plaques

To identify relevant mediators and effectors of fibrous cap stability, we performed laser capture microdissection of fibrous caps in atherosclerotic plaques (10 stable *vs*. 10 unstable plaques) from patients undergoing carotid endarterectomy (CEA) in our Department for Vascular and Endovascular Surgery at the Klinikum rechts der Isar of the Technical University Munich, Germany. Here, *NUDT6* was enriched while *FGF2* expression was downregulated in unstable/ruptured compared to stable fibrous caps. Plaque stability was assessed based on fibrous cap thickness (>/< 200 µm) as previously described (17,26,27) (Figure 1A, B and C). *NUDT6* and *FGF2* form a prototypical mRNA-NAT sense/antisense (S/AS) pair with overlapping exons (Supplemental Figure 1A) previously described in the cancer field (28). *In situ* hybridization (ISH) on advanced atherosclerotic carotid plaques confirmed the increase in *NUDT6* signal predominantly associated with the fibrous cap while showing higher expression in ruptured lesions (Figure 1D). In the same lesions, deregulation of FGF2 levels (high in stable, low in ruptured plaques) assessed by immunohistochemistry (IHC) could be detected (Figure 1E). Smooth muscle cell α-actin (αSMA), an indicator of SMC content and fibrous cap stability, was positively correlated to FGF2 levels and appeared as expected higher in stable and lower in unstable/ruptured lesions (Supplemental Figure 1B). Quantitative real-time PCR (qRT-PCR)-based expression analysis for *NUDT6* and *FGF2* on bulk lesions comparing stable *vs*. plaque phenotypes showed an increase of both *NUDT6* and *FGF2* (Figure 1, F and G) in ruptured lesions. Thus, the deregulation of both transcripts seemed to appear exclusively in the culprit fibrous cap area and not in other regions of plaques, such as the necrotic core or the remaining media of the carotid arteries (Figure 1A).

**Figure 1:**
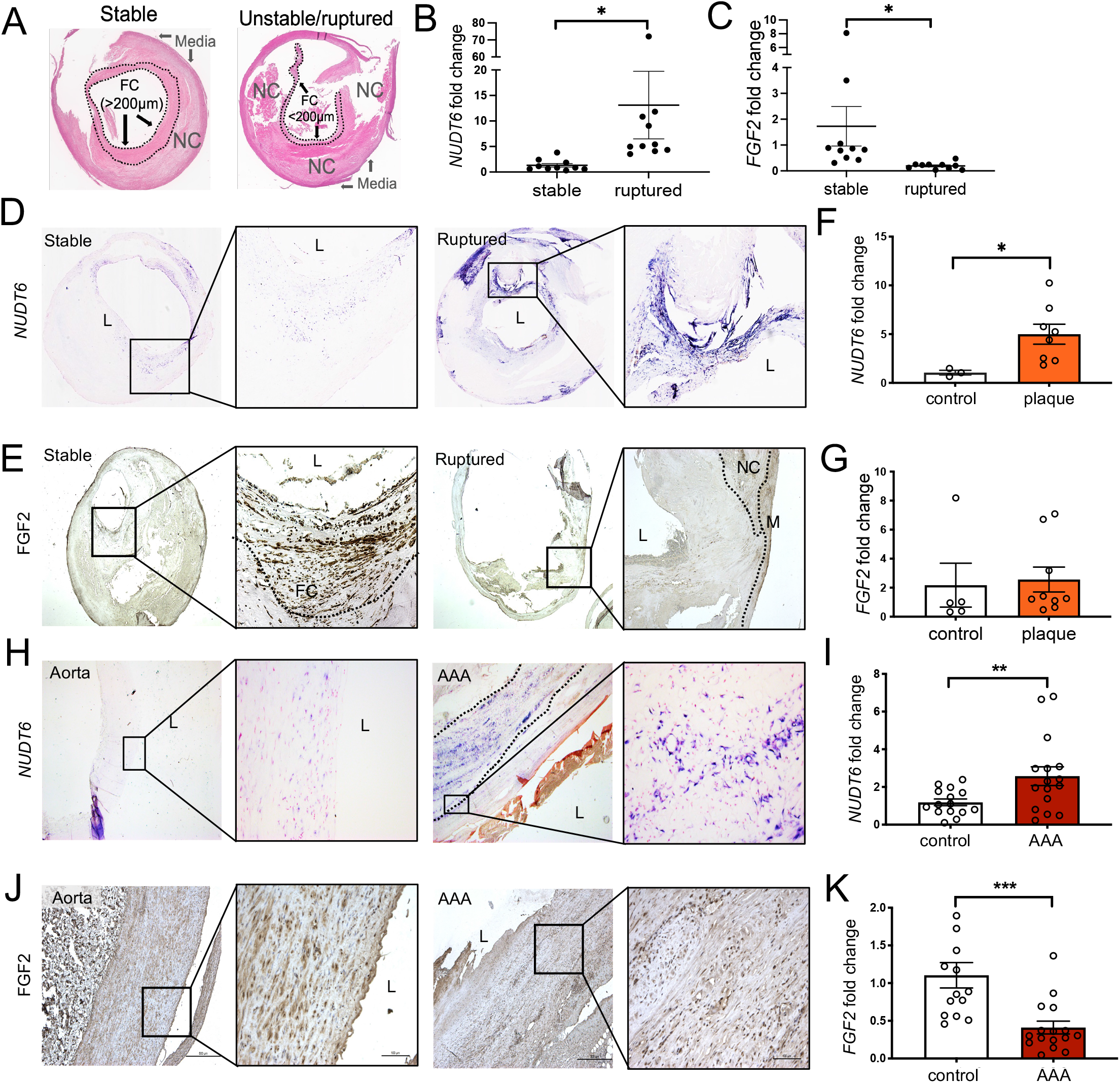
*NUDT6* expression is upregulated in human ruptured carotid plaques and human abdominal aortic aneurysm. **(A)** Haematoxylin and Eosin stained stable and unstable or ruptured atherosclerotic plaques from the Munich Vascular Biobank. The dotted lines mark the Fibrous Cap (FC) shielding the necrotic core (NC) from the lumen. FC>200µm defines stable lesions, FC<200µl defines unstable lesions. **(B-C)** *NUDT6* **(B)** and *FGF2* **(C)** expression in micro-dissected carotid fibrous caps of ruptured *vs*. stable lesions (n=10 per group). **(D)** *In situ* hybridization (ISH) of stable and ruptured atherosclerotic carotid lesions (n=5 per group) indicates a stronger signal in the fibrous caps of ruptured lesions. L= Lumen. **(E)** Immunohistochemical analysis of FGF2 shows decreased expression in correlation to *NUDT6* in fibrous cap ruptured lesions compared to stable lesions (n=3 per group). Dashed lines mark the fibrous cap. L= lumen, FC= fibrous cap, M=media, NC=necrotic core. **(F-G)** qRT-PCR analysis detecting *NUDT6* **(F)** and *FGF2* **(G)** in whole fresh frozen carotid arteries of controls (n=3-5) or advanced lesions (n=8). **(H)** *In situ* hybridization of control aorta and abdominal aortic aneurysm (AAA) (n= 3 per group) indicates higher expression of *NUDT6* in the vessel’s smooth muscle cell-rich media layer (dashed lines). L=lumen. **(J)** Immunohistochemical staining of FGF2 shows decreased expression of both markers in AAA compared to aortic control (n=3 per group). L=lumen. **(I-K)** qRT-PCR of whole fresh frozen abdominal aortic aneurysm (n=16) or control aorta (n=13) detecting *NUDT6* **(I)** and *FGF2* **(K)**. n=13-16. Quantitative data are presented as mean + SEM. *P<0.05; **P<0.01; ****P<0.0001. Significance was determined using one-tailed Student’s t-test **(B-C; F-G; I-K)**.

### *NUDT6* and *FGF2* deregulation is also present in human tissue specimens from patients with abdominal aortic aneurysms

Based on the findings in the fibrous cap from stable and unstable plaques, we wanted to assess whether the deregulation of the *NUDT6*/*FGF2* axis is only present in carotid artery disease or potentially also of relevance in other vascular diseases, in which SMC survival appears of great importance. As this is the case in AAAs, we monitored the expression of both transcripts by qRT-PCR and ISH in human specimens from patients undergoing elective open aneurysm repair. *NUDT6* expression patterns reflected those observed in fibrous caps, with significantly higher AAA levels than non-dilated control aortas (Figure 1,H and I). In line with the fibrous cap results, FGF2 levels were substantially decreased in human AAA on both the protein and RNA levels (Figure 1, J, and K). Also, αSMA, being mainly expressed by contractile SMCs in AAA tissue, was less prominent in AAA *vs*. controls (Supplemental Figure 1C).

### *Nudt6* is increased, and *Fgf2* downregulated in experimental murine models of inducible plaque rupture and abdominal aortic aneurysm development

As both *NUDT6* and *FGF2* are conserved in vertebrates (29), we addressed whether similar disease-associated expression patterns could be observed in mouse models of vascular disease development. Atherosclerotic *ApoE*-deficient mice were utilized for the combined incomplete ligation/cuff injury model we established previously (17,26,27) to induce plaque rupture in carotid arteries (Figure 2A), or the Angiotensin II (AngII) model (30) to create suprarenal aortic aneurysms (Figure 2B). *Nudt6* transcript and Fgf2 protein distribution within the vasculature were visualized *via* ISH and IHC, respectively. In line with our observations in human tissue, *Nudt6* was increased in both disease models (Figure 2C and D), whereas Fgf2 protein was decreased in ANGII-induced aneurysms and ruptured murine plaques (Figure 2E and F). To further mimic our efforts in analyzing advanced human carotid artery lesions, we performed qRT-PCR analysis from microdissected laser-captured murine fibrous caps. This confirmed *Nudt6* upregulation and *Fgf2* mRNA reduction in the cap (Figures 2G and 2H). Similar expression patterns were obtained from qRT-PCR analysis of *Nudt6* and *Fgf2* in AngII-AAA tissue *vs*. saline-infused control specimens (Figures 2I and J).

**Figure 2:**
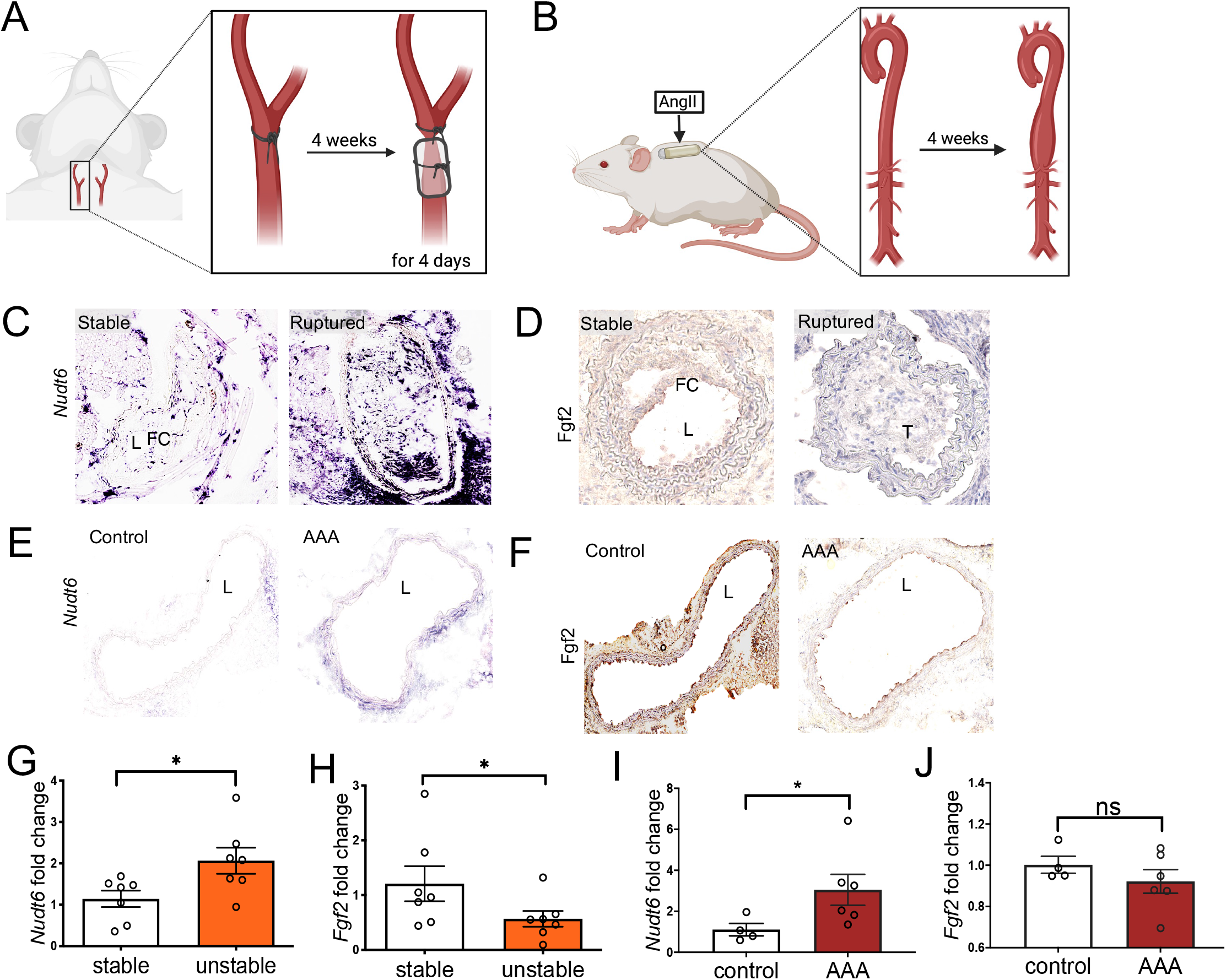
Increased *Nudt6* mRNA and decreased *Fgf2* mRNA and protein levels in two experimental animal models for vascular disease. **(A)** Scheme describing the inducible plaque rupture mouse model. A ligation is placed distally to the carotid bifurcation. After 4 weeks, a cone-shaped cuff is placed around the vessel, leading to disturbed blood flow. **(B)** Scheme of the Angiotensin II mouse model. After the implantation of an AngII-filled pump, an abdominal aortic aneurysm develops. **(C-D)** Stable *vs*. ruptured carotid plaques derived from the murine inducible plaque rupture model (n=7 per group) were stained for *Nudt6* (**C**, *in situ* hybridization) and Fgf2 (**D**, immunohistochemistry), resembling the phenotype observed in human disease with increased *Nudt6* mRNA expression and decreased Fgf2 protein levels in ruptured compared to stable lesions. (n=3 per group). **(E – F)** Saline-infused (control) *vs*. AngII-infused (AAA) aorta from the Angiotensin II mouse model (n=6 per group) was stained for *Nudt6* **(E)** and Fgf2 **(F). (G-J)** The observed phenotype was confirmed by qRT-PCR in murine stable *vs*. unstable/ruptured lesions **(G-H)** and murine saline-infused (control) *vs*. AngII-infused aortas **(I-J)** (n=4-7 per group). Quantitative statistics are presented as Mean+SEM. *p<0.05 in one-tailed Student’s t-test.

### *Nudt6* inhibition *in vivo* reduces rupture risk and AAA growth

To test the therapeutic feasibility of inhibiting *Nudt6 in vivo*, locked nucleic acid (LNA) GapmeRs were delivered to the above-mentioned experimental mouse models. The designed GapmeR targeted *Nudt6* at exon 2, which does not overlap with *Fgf2* (see Supplemental Figure 1A). Inhibiting *Nudt6* in the inducible plaque rupture model led to decreased rupture rates (Figure 3A). ISH and IHC further confirmed reduced *Nudt6* expression while rescuing Fgf2 and αSma expression in GapmeR-treated murine carotid arteries (Figure 3B, Supplemental Figure 1D). Next, local inhibition of *Nudt6 via* ultrasound-targeted microbubble destruction (UTMD) was carried out in the aorta of AngII-treated mice with AAA. As shown in a previous study (27), local delivery of antisense oligonucleotides (ASOs) can limit potential off-target effects while potentiating the inhibitory action of the site-specific GapmeRs applied in our experiments. Local penetration of the aortic tissue was confirmed by sectioning and imaging of the abdominal aorta treated with a fluorescent FAM-labelled probe (Supplemental Figure 1E). *Nudt6*-Antisense Oligonucleotide (ASO, GapmeR) treatment resulted in decreased abdominal aortic diameters upon weekly treatments until day 28 (Figure 3C). Furthermore, an overall improved vessel wall morphology (intactness of elastic laminae, increased SMC content) was accompanied by decreased *Nudt6*, as well as enhanced Fgf2 and αSma expression levels (Figure 3D, Supplemental Figure 1D-F).

**Figure 3:**
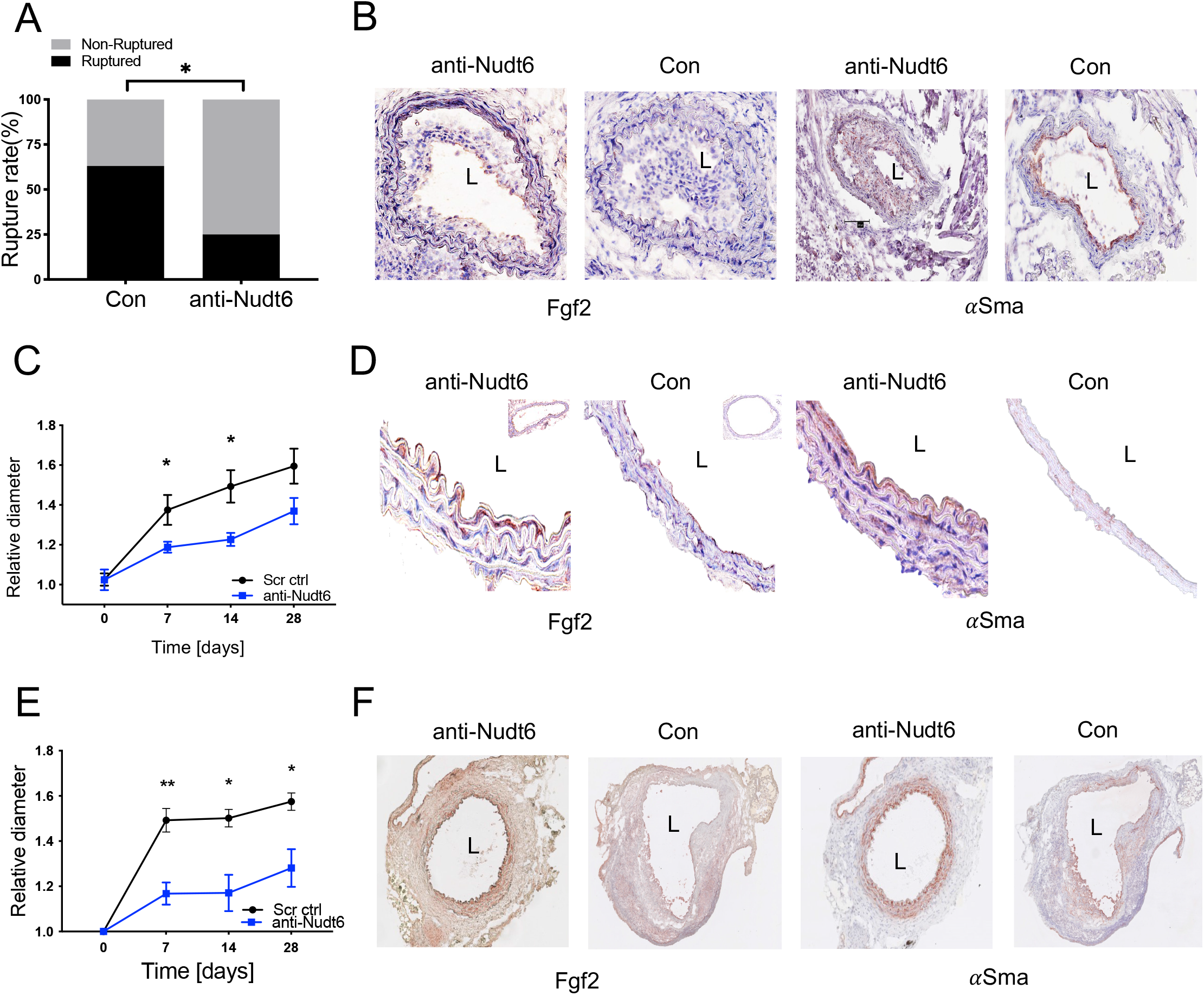
Modulating *Nudt6 in vivo* leads to reduced rupture risk and reduced AAA growth. **(A)** *Nudt6*-ASO treatment (n=20) of *ApoE* ^-/-^ mice significantly reduced the rupture rate in the inducible plaque rupture model compared to scramble-control treated animals (n=20). **(B)** Fgf2 and *α*Sma immunohistochemistry display restored expression after *Nudt6*-ASO treatment (n=5 per group). **(C)** In the AngII model, local *Nudt6*-ASO treatment (n=8) *via* ultrasound-targeted microbubble destruction (UTMD) led to significantly lower abdominal aortic diameter and reduced growth compared to scramble control treatment (n=13). **(D)** Fgf2 and *α*Sma protein levels are restored in *Nudt6*-ASO treated animals (n=3 per group). **(E)** Systemic *Nudt6*-ASO treatment (n=4) in the porcine pancreatic elastase (PPE) mouse model significantly reduced abdominal aortic diameter and growth rate compared to control (n=5). **(F)** Immunohistochemistry of Fgf2 and *α*Sma indicates higher expression in the *Nudt6*-ASO group compared to the control group (n=3 per group). Quantitative data are shown as mean + SEM. *p<0.05, **p<0.01. Significance was calculated using χ2 test **(A)**, and multiple t-tests **(C, D)**.

To further confirm the observed findings of the AngII-AAA model, we performed a confirmatory study in a second mouse model of AAA development using porcine pancreatic elastase (PPE) in C57BL/6 mice. Utilization of this model allowed us to discriminate potential model-specific effects of AngII-induced aneurysms on *Nudt6* and *Fgf2* deregulation. A one-time systemic intraperitoneal injection of the *Nudt6*-ASO was performed on the same day the AAA was induced. Similar to our observation in AngII-treated mice, downregulation of *Nudt6* (Supplemental Figure 1D) resulted in decreased abdominal aortic diameters (Figure 3E). Notably, Fgf2 and αSma content within the *intima-media* was increased in the *Nudt6*-ASO group compared to scrambled GapmeR control mice with AAAs (Figure 3F). To conclude, *Nudt6* inhibition in three different mouse models led to alleviated and beneficial effects, such as reduced rupture rate of fibrous caps and AAAs, as well as smaller aortic diameters compared to respective control groups.

### *NUDT6* inhibition derepresses FGF2, resulting in increased proliferation and reduced apoptosis in smooth muscle cells

We next investigated *NUDT6*-mediated FGF2 regulation by using different stimuli *in vitro* relevant to vascular disease development and progression. Oxidized low-density lipoprotein (oxLDL) triggers foam cell formation during atherogenesis in macrophages and SMCs (31). As we identified *NUDT6* being enriched in SMCs, we stimulated human carotid smooth muscle cells (hCtSMCs) with oxLDL and/or serum starvation. *NUDT6* RNA levels increased upon oxLDL stimulation, whereas *FGF2* was strongly reduced dose-dependent (Figure 4A). Similarly, human aortic smooth muscle cells (hAoSMCs) were treated with AngII, a factor known to trigger vascular inflammation and AAA formation (32). AngII stimulation resulted in greatly increased *NUDT6* expression while reducing *FGF2* levels (Figure 4B). Furthermore, primary aortic SMCs isolated from three different AAA patients, as previously described (33) showed a significant upregulation of *NUDT6* with a concomitant reduction of *FGF2* levels compared to commercially available hAoSMCs (Supplemental Figure 2A and B). This could be reversed by knocking down *NUDT6 via* siRNAs (Supplemental Figure 2C and D). As *NUDT6* expression seemed to be triggered by disease, we subsequently tested the effects of *NUDT6* knockdown (KD) and overexpression (OE) in hCtSMCs and hAoSMCs. siRNA-mediated silencing of *NUDT6* (Supplemental Figure 2E) caused significant FGF2 upregulation on both mRNA (Figure 4, C and E) and protein levels (Figure 4D). Conversely, *NUDT6* OE substantially decreased *FGF2* mRNA (Figure 4C) and protein levels (Figure 4F).

**Figure 4:**
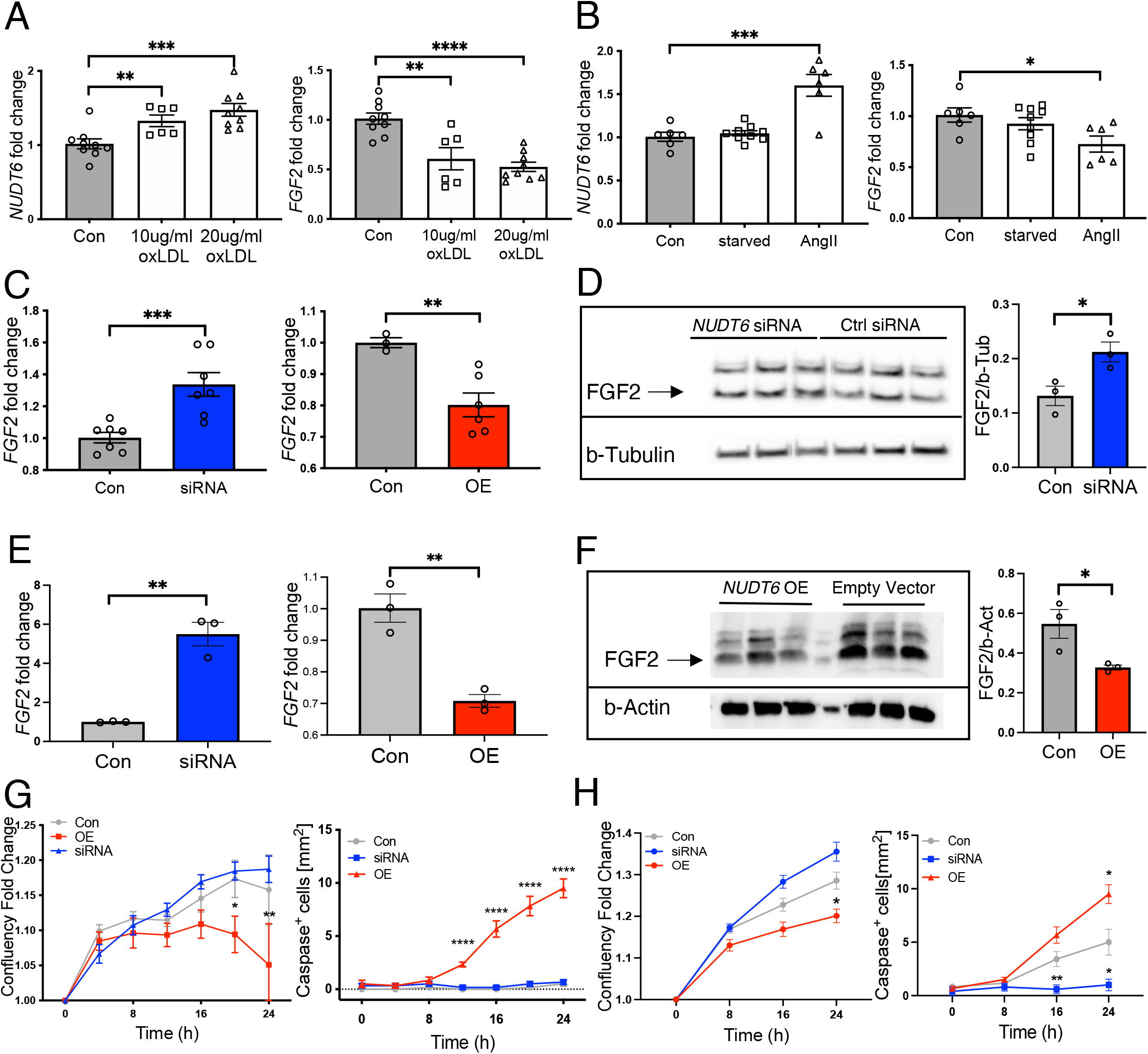
*NUDT6* represses *FGF2*, thereby inducing apoptosis and inhibiting proliferation. **(A)** Oxidized LDL treatment results in a dose-dependent increase of *NUDT6* expression, whereas *FGF2* decreases with increased dosage in human carotid smooth muscle cells (hCtSMCs) (n=6 per group). **(B)** Angiotensin II treatment of human aortic smooth muscle cells (hAoSMCs) results in increased *NUDT6* and decreased *FGF2* expression (n=6-9 per group). **(C-E)** By transfection of hCtSMCs **(C)** and hAoSMCS **(E)**, siRNA-mediated knockdown of *NUDT6* results in increased *FGF2* expression, whereas vector-mediated overexpression of *NUDT6* limits *FGF2* mRNA levels. N=6 per group. **(D)** *NUDT6* silencing restores FGF2 protein, n=3 per group. **(F)** *NUDT6* overexpression leads to less FGF2 protein expression. **(G-H)** Dynamic live-cell imaging of both hCtSMCs **(G)** and hAoSMCs **(H)** show impaired proliferation capacity in *NUDT6* overexpressing cells while the apoptosis rate is significantly increased. *NUDT6*-siRNA treated cells behave as scramble control-treated cells. N≥6. Quantitative results are presented as mean + SEM. *p<0.05; **p<0.01; ***p<0.001; ****p<0.0001. Significance is determined using one-way ANOVA with Sidak **(A-B)**, One-tailed Student’s t-test **(C-F)** or two-way ANOVA with Tukey **(G-H)**.

By using a dynamic live cell imaging system, we further explored the effects of *NUDT6* KD and OE, respectively, in terms of proliferation and apoptosis rates. Silencing of *NUDT6* did not impair proliferation or apoptosis in either hCtSMCs or hAoSMCs (Figure 4G and H). However, overexpression of *NUDT6* in both cell types limited proliferation and increased apoptosis. Migration and motility of SMCs were also worsened in cells with *NUDT6* OE, but not in cells where *NUDT6* was silenced (Supplemental Figures, 3F and G).

In addition to its regulatory function as an antisense transcript, a micropeptide coding function for *NUDT6* was recently described (34,35). To rule out any functional effect of the *NUDT6* micropeptide on SMCs, we stimulated hCtSMCs with the *NUDT6* peptide (Supplemental Figure 2H). Importantly, we were not able to detect any functional role for this micropeptide in SMCs, supporting the fact that *NUDT6* RNA seems solely relevant for vascular disease development and progression.

To determine where *NUDT6* and *FGF2* are located in the cell, nuclear-cytoplasmic fractionation was performed. This revealed localization of both *NUDT6* and *FGF2* primarily in the nucleus and to a lesser extent in the cytoplasm of hAoSMCs (Supplemental Figure 3I). In hCtSMCs, *NUDT6* was found to be equally distributed between cytoplasm and nucleus (Supplemental Figure 2J). Globally, these results suggest a worsened outcome in terms of aortic and carotid SMCs survival and migration upon *NUDT6* upregulation, as induced by pathogenic stimuli, such as oxLDL and AngII, or by *NUDT6* overexpression. The ability of hCtSMCs to shield the necrotic core is impaired, while hAoSMCs fail to maintain the aortic wall integrity, leading to rapid expansion and acute rupture.

### *NUDT6* interacts with CSRP1 and thereby affects smooth muscle cell contractility and migration

To further explore the distinct molecular mechanism underlying *NUDT6* regulation, we aimed to identify a potential mediating function through direct interaction with other proteins. We thus performed a *NUDT6-*RNA pulldown experiment followed by liquid chromatography-mass spectrometry (LC-MS/MS). Using hAoSMC lysate and *in vitro* transcribed biotinylated *NUDT6* or control, 50 targets were identified as significantly enriched in the *NUDT6* pulldown fraction (Figure 5A). Cysteine and Glycine Rich Protein 1 (CSRP1, 32.6-fold increase, Supplemental Table 1) appeared most enriched and was thus chosen for further in-depth analysis based on its previously reported relevance to SMC function (Supplemental Figure 3A) (36-38). In general, many interaction partners seemed to be involved in SMC-mediated vascular pathologies (Supplemental Figures 3B and C). The direct *NUDT6*-CSRP1 interaction was further confirmed using RNA Immunoprecipitation (RIP), where *NUDT6* could be detected in the eluted CSRP1 fraction (Figure 5B).

**Figure 5:**
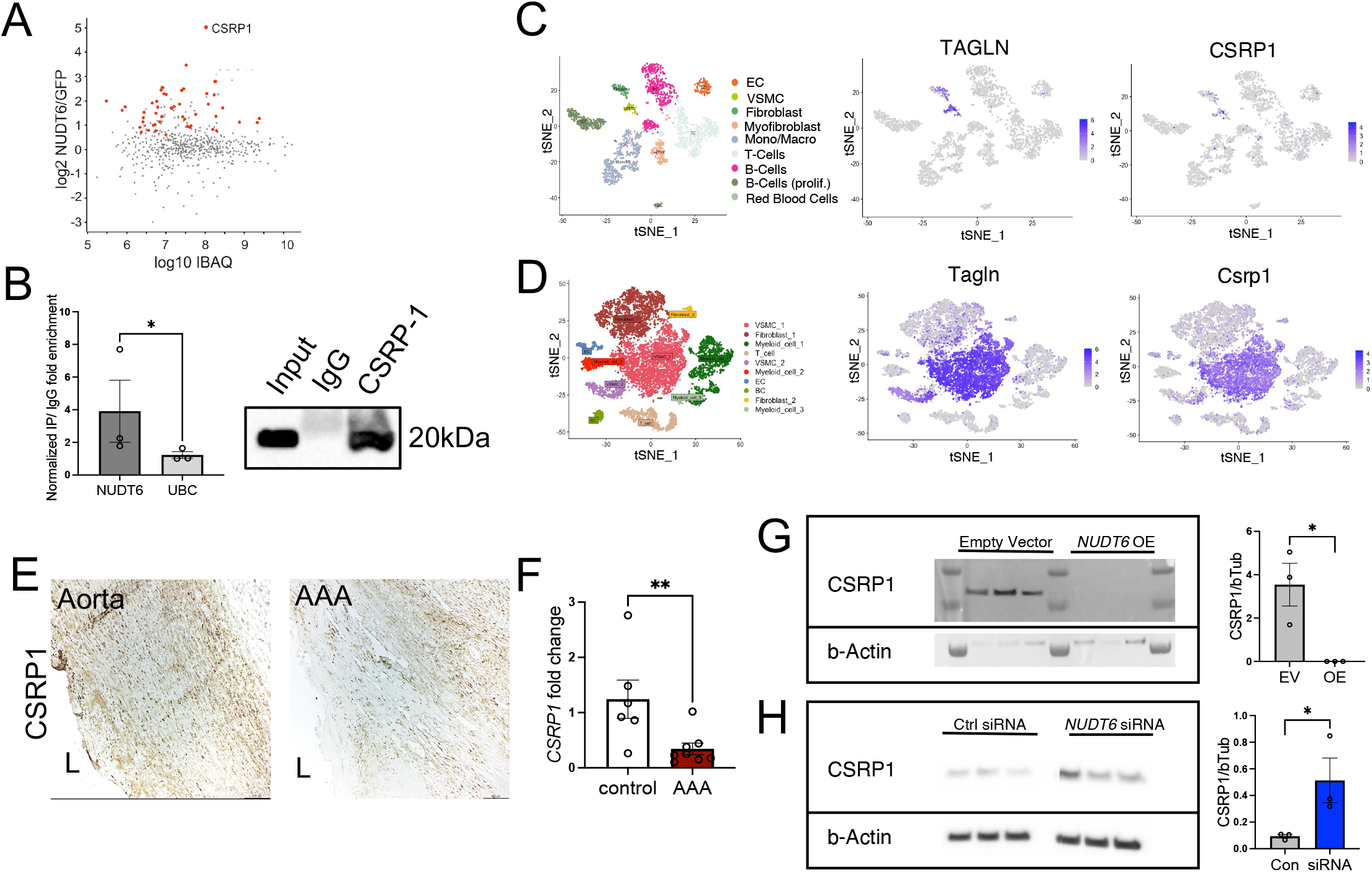
Identification and Validation of NUDT6 interaction partner CSRP1. **(A)** Identified proteins in biotinylated NUDT6 versus biotinylated GFP pulldown in hAoSMCs (n=6 per condition), red dots represent significantly enriched proteins (Student’s t-test) to NUDT6. **(B)** Confirmed binding of NUDT6 to CSRP1 protein in hAoSMCs. NUDT6 and UBC (unrelated target) enrichment in CSRP1 IP fraction was quantified using RTqPCR and CSRP1 availability was verified via Western Blot. T-distributed stochastic neighbor embedding (t-SNE) plots of single cell RNA sequencing data from human **(C)** human and **(D)** murine AAA (PPE-induced) with featured expression plots of Transgelin (TAGLN) and CSRP1. **(E)** Immunohistochemical staining of CSRP1 in human control aorta and AAA. (L’ indicates lumen). **(F)** qRT-PCR of whole fresh frozen abdominal aortic control (n=6) and AAA (n=8) of CSRP1 mRNA. **(G)** CSRP1 protein levels after NUDT6 overexpression. **(H)** CSRP1 protein levels after NUDT6 siRNA-mediated knockdown (n=3 per group). Quantitative data are shown as mean + sem *p<0.05; **p<0.01. Significance was calculated using one-tailed Student’s t-test. EC= Endothelial Cells, VSMCs = Vascular Smooth Muscle Cells; Mono/Macro=Monocytes/Macrophages; B-Cells (prolif.) = proliferating B-Cells

Next, we performed single-cell RNA sequencing (scRNA-seq) on human AAA tissues, as well as murine and porcine PPE-induced AAAs to gain insight into the specific cellular expression pattern of our targets. Of importance, *CSRP1* expression seemed mainly restricted to SMCs (Figure 5C and D, Supplemental Figure 3D) in all three species. This observation was confirmed by IHC staining of human control aorta and AAA, where CSRP1 levels were reduced in AAA compared to healthy aortic tissue (Figure 5E). Further, significantly lower *CSRP1* mRNA levels were found in AAA (Figure 5F). Of note, both *NUDT6* and *FGF2* were discovered but lowly expressed in the human and porcine scRNA-Seq datasets. However, the two transcripts were found to be higher expressed in murine AAAs (Supplemental Figure 3E). To further study the effect of *NUDT6* on CSRP1, we modulated its expression in hAoSMCs using plasmid-based overexpression and GapmeR-mediated knockdown. Overexpression of *NUDT6* substantially downregulated CSRP1 (Figure 5G), while inhibition of *NUDT6* increased its protein levels (Figure 5H). This indicates the regulatory function of *NUDT6* on SMC migration, survival, and de-differentiation *via* targeting of CSRP1 (36,38). This effect is independent, but additive in vascular disease development, to its antisense regulation of the *FGF2* gene.

### *NUDT6* inhibition limits aortic aneurysm development in *LDLR-*deficient Yucatan mini-pigs

Finally, we attempted a translational therapeutic intervention in a novel preclinical large animal model of AAA disease, utilizing *LDLR*-deficient Yucatan mini-pigs. One-year-old male *LDLR*^-^ _/-_mini-pigs were fed a high-cholesterol Western diet for six months to trigger advanced atherosclerotic disease (Supplemental Figure 4A). AAAs for this study were induced *via* catheter-directed PPE infusion into the infrarenal section of the porcine aortas as previously described (32,33). Site-specific GapmeRs targeting porcine *NUDT6* were tested in pig fibroblasts before application (Supplemental Figure 4B) to ensure sufficient knockdown. To potentiate efficient *NUDT6* inhibition and enhance our study’s translational feasibility, a local application using drug-eluting balloons (DEBs) coated with *NUDT6*-ASO was chosen. Seven days after AAA induction (mimicking a small but clinically relevant aneurysm in humans), *NUDT6*-ASO were locally applied to target the AAAs *via* one-time endovascular DEB treatment. A sham intervention was performed using vehicle-only coated DEBs and served as the control group.

From the time of intervention, we realized a significant difference in the relative aortic diameters between the two groups. *NUDT6*-ASO DEB-treated animals displayed significantly smaller diameters than the control group (Figure 6C). Profound changes in *FGF2* and *ACTA2*/αSMA expression on protein and mRNA levels could be observed in the treatment *vs*. the control group (Figure 6D and E). These results were further confirmed on the mRNA level by qRT-PCR (Figure 6F-G, Supplemental Figure 4C). In addition, expression profiles of the other direct *NUDT6* RNA-protein interaction partner, CSRP1, resembled our *in vitro* results, with increased expression in the aortas from pigs receiving the *NUDT6*-ASO treatment (Figure 6D). The overall aortic morphology, including elastic layer integrity, SMC shape (not spindle-like), and content (yellow staining in Elastica van Giesson indicates SMCs), immune cell infiltrates, and atherosclerotic plaque burden was less altered compared to internal control tissue specimens (Supplemental Figure 4D).

**Figure 6:**
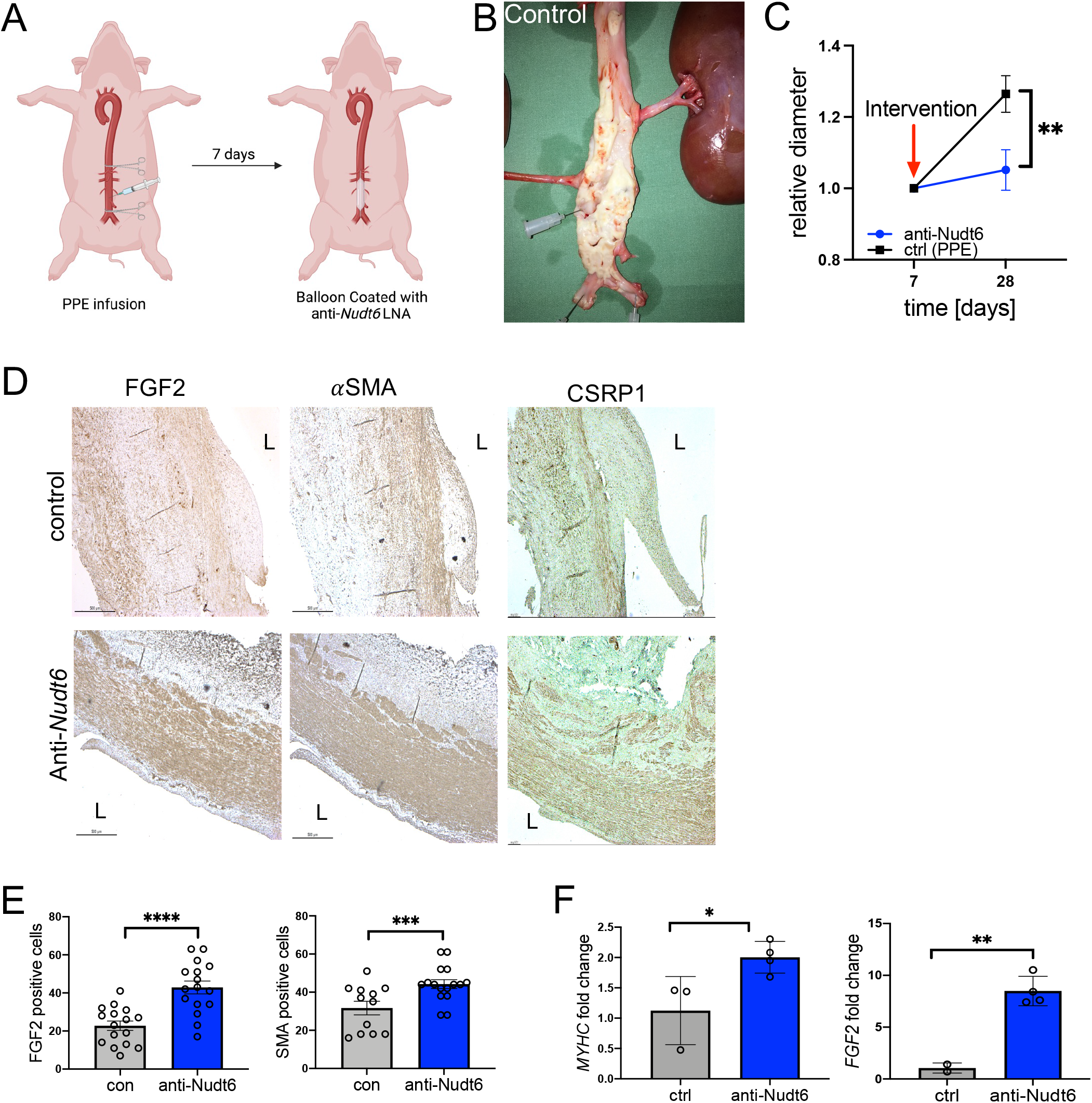
Translational aspects of *NUDT6* inhibition in a large animal model of AAA. **(A)** Scheme describing the AAA induction in *LDLR*^-/-^ Yucatan minipigs, in which the abdominal aorta was clamped and infused with PPE for 1 minute. 7 days later, a balloon coated with *NUDT6*-ASO was introduced *via* the femoral artery and inflated for 3 minutes. **(B)** Representative image of the abdominal aorta of a control *LDLR*^-/-^ minipig. **(C)** The relative diameter of *NUDT6*-ASO-treated animals (n=4) shows a decrease compared to DEB control-treated animals (n=3). **(D)** Immunohistochemical staining of the abdominal aorta of control and *NUDT6*-ASO receiving pigs using antibodies against FGF2, *α*SMA, and CSRP1. “L” indicates lumen. **(E)** Quantification (4 high power fields per aorta, in total 16 counts) of immunohistochemical staining from **D** for FGF2 and *α*SMA. (**F-G)** qRT-PCR of whole porcine aortic tissue shows a significant increase in *MYHC* **(F)** and *FGF2* **(G)** mRNA expression in *NUDT6*-ASO-treated animals (n=4) compared to control (n=3). Quantitative results are presented as mean + SEM. *p<0.05, **p<0.01, ***p<0.001; ****p<0.0001. Significance was determined using two-way ANOVA with Sidak **(C)** and one-tailed Student’s t-tests **(E-G)**.

## Discussion

Increasing evidence shows that several non-coding RNA subclasses (*e*.*g*., microRNAs, lncRNAs) play an essential role in mediating gene expression in health and disease. LncRNAs regulate gene expression transcriptionally as well as post-transcriptionally by affecting the stability and translation of protein-coding mRNAs. The numbers of ncRNAs (and possibly their functions) have increased with evolution, perhaps explaining how organisms developed complexity without a corresponding increase in protein-coding genes (39). However, it is still being determined how many of these transcripts, particularly those with low abundance, are functional or transcriptional artifacts. Nonetheless, the number of annotated ncRNAs in humans continues to grow and has already surpassed the amount of protein-coding genes by far.

The explosion of new RNA classes has raised the possibility of new therapeutic strategies that mimic or antagonize their function. Attempts to therapeutically target RNA pathways began as soon as they were discovered, even before their mechanism of action had been revealed. On the contrary, within the roughly 20.000 expressed human proteins, only a fraction of 10–15% are thought to be involved in disease development and exacerbation (40). Thus, disrupting their activity is likely to have a therapeutic benefit in patients with cardiovascular disease. Most potentially disease-related proteins are termed ‘undruggable’, implying that they lack distinctive cleft-like motifs into which small molecules can bind with high affinity and specificity (41). These ‘undruggable’ proteins contain many transcription and growth factors, as well as epigenetic mediators. In this dilemma, several RNA-based drugs have shown clinical benefits for treating patients with previously considered intractable diseases (42). Most novel drugs that interfere with RNA expression use nucleic acid analogs, taking advantage of complementary base-pairing to mimic or antagonize endogenous RNA processes like the GapmeRs (ASOs) utilized in our *in vivo* experiments.

In our present study, we report a substantial imbalance of *NUDT6* and its sense target FGF2 in the context of two different vascular diseases: advanced atherosclerosis and abdominal aortic aneurysms. Both diseases share common key features, such as tissue infiltration of inflammatory cells, impaired ECM remodeling, and SMC apoptosis. The latter process is known to further be associated with the thinning of fibrous caps and larger necrotic cores (43) in carotid artery plaques, while in AAA, SMC apoptosis has been extensively acknowledged as a major process underlying aneurysm formation and progression. In particular, upregulation of *NUDT6* and a concomitant downregulation of *FGF2* mRNA and protein were observed within fibrous caps of vulnerable atherosclerotic human plaques and in aneurysmal aortas from patients undergoing elective open repair surgery. A significant focus of our study was to identify molecular circuits and to develop novel therapies aimed at halting late-stage disease events, such as the destabilization and vulnerability of an advanced plaque and acute rupture of AAA.

One possibility to prevent these events is stabilizing the vessel wall or fibrous cap by re-activating proliferation and migration pathways in de-differentiated/senescent SMCs. We were able to show that de-repression of FGF2 *via* inhibiting *NUDT6* in both carotid and aortic SMCs, led to increased proliferation, migration, and decreased apoptosis. In contrast, the opposite could be observed in gain-of-function studies using *NUDT6* overexpression. Upon *NUDT6* targeting, SMCs regained their migratory capacity and contributed to forming a thicker fibrous cap in carotid atherosclerosis. The same was true in dilated aortas, where *NUDT6* inhibition strengthened the weakened vessel wall. The identified protein interaction partner of *NUDT6*, CSRP1, further points towards a dynamic regulatory role of *NUDT6* in SMCs in health and disease states apart from acting as an antisense to *FGF2*. CSRP1 is a co-activator of the serum response factor (SRF), which controls SMC differentiation and proliferation (36-38).

It was shown previously that applying basic Fibroblast Growth Factor (bFGF/FGF2) to the abdominal aorta *via* a biodegradable hydrogel halted aneurysm progression and restored the contractile capacity of SMCs (44,45). Also, in intracranial aneurysms, the lncRNA *ANRIL* was shown to influence the proliferation of SMCs by positively regulating FGF2 (46). However, the role of FGF2 in atherosclerosis is more complex and dynamic: in early stages, it can support macrophage infiltration (47) while triggering intimal hyperplasia and thickening (48). It has however also been shown that in combination with PDGF-BB, high FGF2 levels can be beneficial by structurally normalizing abnormal neovessels and thereby improving plaque stability (49). This is in line with our observations where *FGF2* mRNA expression in microdissected fibrous caps was strongly deregulated. In contrast, *FGF2* mRNA from whole carotid plaque tissues did not differ in expression between stable and unstable/ruptured lesions. Thus, it seems to be dependent on the progression of the respective disease to determine when to re-activate FGF2, as well as on the location of the FGF2 signal within the advanced plaque (media, necrotic core, or fibrous cap).

Using GapmeRs targeting *NUDT6* in four different animal models from two different species, we could trigger FGF2 at a late time point of the disease and observed positive effects on rupture rate and abdominal aortic diameter, respectively. Systemic injection of *NUDT6* GapmeR in the murine inducible plaque rupture model and the murine PPE-induced AAA model led to different effects on *NUDT6* mRNA, as well as FGF2 and αSMA protein and mRNA levels. Based on this, the overall composition of the vasculature appeared less diseased and better remodeled. In the murine AngII and the porcine PPE AAA models, we applied *NUDT6* GapmeR locally to the aortic wall using two different methods, which both increase the translational value of utilizing *NUDT6* GapmeRs. UTMD, successfully used for clinical trials in patients with inoperable pancreatic cancer (50), appears as a feasible and translational approach for delivering ASOs to the abdominal aorta. Interestingly, in patients with pancreatic cancers, the previously deemed ‘undruggable’ transcript *KRAS* was targeted by employing local RNA therapy utilizing a polymeric matrix containing siRNA (51,52). These findings pinpoint the necessity and opportunity of potent local administration of RNA therapeutics, such as siRNAs, ASOs, and antagomiRs.

Based on the idea of enhanced local delivery, we utilized DEBs in male and female *LDLR*^-/-^ Yucatan minipigs in a preclinical AAA model, demonstrating the feasibility of enhanced *NUDT6* inhibition. This is the first study in the cardiovascular field to target a lncRNA with ASOs in a large porcine animal model (53). MicroRNAs have been successfully targeted before in large animal models (53) and are currently being assessed in phase II clinical trials for treatment in patients with myocardial ischemia (54) and heart failure (55). We are convinced that our study provides evidence and hope that inhibition of *NUDT6* presents a novel therapy to limit the burden of atherosclerotic vascular diseases, such as advanced carotid stenosis and AAAs. Future studies will aim at optimizing the local delivery of the *NUDT6*-ASO strategy into carotid plaques and aortic aneurysms to successfully treat our increasing number of patients with both vascular pathologies.

## Material and Methods

### Laser-capture microdissection of fibrous cap tissue derived from advanced atherosclerotic carotid plaques

Human specimens were treated as described in the Supplemental Methods. Murine specimens were embedded in OCT medium (Sakura Fine-Tek, Umkirch, Germany) and stored at −80°C until further processing. Before H&E staining, mouse sections were fixed in 4%PFA (Sigma-Aldrich, Darmstadt, Germany) for 10min. H&E was performed on RNase-free glass slides and pre-treated with UV light at 254nm for 30 min to promote better adhesion to the slide. Up to 10 consecutive slides per patient were micro-dissected and collected in 350µl of RLT buffer (Qiagen). The sample was inverted and centrifuged at 13.000rpm for 5 min and stored at −80°C. RNA extraction was performed with the RNeasy Micro Kit (Qiagen, Hilden, Germany) after the manufacturer’s instructions. RNA quality was verified using Agilent 2100 Bioanalyzer (Agilent Technologies), and RNA quantity was measured using Nanodrop (Agilent Technologies). For further information, please refer to Jin *et al*.(27).

### *In Situ* Hybridization

Frozen sections were thawed and hybridized with Exiqon miRCURY LNA GapmeR microRNA ISH Optimization Kit (Exiqon, Denmark or Qiagen, Hilden, Germany) following the instructions of the manufacturer with minor modifications (according to Exiqon’s Guidelines for ISH on fresh frozen material using the miRCURY LNA ISH Optimization kits) and using Pertex (VWR, Stockholm, Sweden) for mounting. Slides were stained for *NUDT6* (562759-2, Exiqon, Denmark). Nuclear counterstaining was done with Nuclear Fast Red (Sigma Aldrich, Darmstadt, Germany) for 1 min and then washed in 37°C tap water for 10 min before mounting.

### Mouse carotid plaque rupture model

Via incision of the medial neck, the right carotid artery of an *ApoE*-deficient mouse was cleared from connective tissue. A 5.0 VICRYL suture (Ethicon, Johnson&Johnson AB, Sollentuna, Sweden) was placed around the carotid artery below the bifurcation following the closure of the skin with 5.9 VICRYL suture (Ethicon, Norderstedt, Germany). Twenty-eight days afterward, a 1.77mm long plastic cast consisting of two halves (Promolding BV, Rijswijk, Netherlands) with a triangular-shaped internal lumen ranging from 150µm (distal) to 300µm (proximal) was placed proximal to the ligation around the carotid artery. Before that, flow through the ligated part of the carotid artery was verified by Doppler-enhanced ultrasonography (Vevo 2100, VisualSonics, Toronto, Canada). After the cast was placed, animals were sacrificed, and organs were harvested.

### Mouse Angiotensin II-induced AAA model

Once they reached the age of 10 weeks, osmotic minipumps (model 2004, Alzet, Cupertino, USA) with Ang II (Sigma Aldrich, Stockholm, Sweden) were implanted subcutaneously into the back/neck area of the anesthetized (2.0% Isoflurane-containing oxygenated air) ApoE-deficient mouse on day 1. A steady delivery of 1,000ng/kg/min was thereby ensured. Determination of the aortic diameter *via* ultrasound (VisualSonics; Vevo 2100 Imaging System) was performed at baseline, and then after 7, 14, and 28 days. In addition, the aortic wall was examined for potential dilations and lesions on days 7, 14, and 28. Mice were anesthetized prior to ultrasound with Isoflurane (Santa Cruz Biotechnology, Dallas, USA).

### UTMD-mediated local aortic *Nudt6* GapmeR delivery

The procedure was performed as recently described by Jin et al. (27). In brief, mice subjected to Angiotensin-II osmotic minipumps received 200µl cationic microbubbles (SonoVue), which were charge-coupled to *Nudt6* GapmeR or a FAM-labelled scrambled control (Qiagen, Hilden, Germany). Mice were anesthetized after aortic diameter measurement, and a phased transducer (Sonitron GTS Sonoporation System) was placed on the abdominal aorta, which was marked with a pen according to the localization of the infrarenal aorta. For 15 min, ultrasound was transmitted with 1MHz, 2.0 w/cm^2^, and 50% duty cycles.

### Mouse porcine pancreatic elastase–induced AAA model

The proximal and distal aorta of anesthetized (2.0% Isoflurane-containing oxygenated air) C57BL/6J mice was temporarily ligated at the proximal and distal end. An incision at the bifurcation and an infusion catheter was used for flushing the aorta for 5min at 100mmHg with saline or saline-containing type I PPE (1.5U/ml; Sigma Aldrich, Darmstadt, Germany). Then, the catheter was removed, and the incision was closed with a suture without constricting the aortic lumen. The aortic diameter was measured, as explained in the previous paragraph.

### *In vivo* Nudt6 inhibition

10mg/kg fluorescently labeled scrambled control oligonucleotide or NUDT6 specific GapmeR inhibitor (NUDT6_6 GGACCTGAATTCTGA, Exiqon, Vedbaek, Denmark) were injected intraperitoneally. For the carotid artery atherosclerotic model, mice received a 4-time injection on days 1, 11 and 21, and 28. For the AngII mouse model, mice received 3-time injection on days 1, 11, and 21, followed by the described UTMD procedure. On day 28, the mice were sacrificed through CO_2_ inhalation, exsanguinated by heart puncturing, and perfused with PBS. For the PPE mouse model, mice received a one-time intraperitoneal injection after surgery on the same day. Organs were snap-frozen in dry ice and stored at −80°C. The aneurysmal aorta and the common right carotid artery were embedded into OCT (optimum cutting temperature compound) (Histolab, Gothenburg, Sweden), snap frozen, and stored at −80°C for sectioning.

### Cell proliferation, migration, and apoptosis assay using IncuCyte ZOOM

Minimum 6-8 hours after transfection, cells were live imaged with the IncuCyte imaging system (IncuCyte, Essen Bioscience, Hertfordshire, UK). The experiment was conducted for 24-72 hours, repeated at least two times with triplicates of each group for statistics. To identify the effect of *Nudt6* modulation and its peptides on human carotid smooth muscle cells and human aortic smooth muscles, cell proliferation-(confluence), migration- and apoptosis assays were performed and analyzed by Incucyte® Live Cell Analysis System (Sartorius Stedim Plastics GmbH, Goettingen, Germany) under Nudt6 modulation (overexpression or silencing). IncuCyte® Caspase-3/7 Apoptosis Assay Reagent (Sartorius Stedim Plastics GmbH, Goettingen, Germany) was used to detect apoptotic cells following the manufacturer’s protocol.

### scRNA-seq data analysis

The scRNA-seq datasets were analyzed with Seurat (version 4.1.0) in R (version 4.1.1) (56,57). CellCycleScoring function (58) in Seurat was applied to calculate the scores of cell cycle phases like S phase and G2M phase. SCTransform normalization workflow (59) was adopted to mitigate possible technically driven or other variations in which mitochondrial genes and cell cycle phase were regressed out. In the human dataset, a total of 3000 cells were used for the downstream analysis. In the pig’s models, 6899 cells passed the quality control and were included for analysis, while 13002 cells were analyzed in the mice models. T-SNE (t-Distributed Stochastic Neighbor Embedding) was used for converting cells into two-dimensional maps. FindAllmarkers function was performed to detect the main features of each cluster with default parameters. And the top expressed genes were mainly used for the consideration of cell type identifications.

### RNA Pulldown

2.5ug Nudt6 Human Tagged ORF Clone (OriGene Technologies, Inc. Rockville, MD, USA) or GFP plasmid respectively were digested with XhoI for 1hr (New England Biolabs, Ipswich, MA, USA) for linearization. Both constructs were run on a 1% agarose gel to check for complete linearization. PCR cleanup kit (Qiagen, Hilden, Germany) was used to purify the DNA, which NanoDrop then measured. Next, T7 polymerase (Promega, Walldorf, Germany), as well as the Roche Biotin RNA labeling kit (Sigma Aldrich, Darmstadt, Germany), was used for in vitro transcription (IVT) of 1ug linearized constructs. After that, TURBO DNase (Invitrogen, Darmstadt, Germany) was used to remove the template from the reaction. QuickSpin Sephadex columns (Roche, Sigma Aldrich, Darmstadt, Germany) were used to remove any unwanted leftovers from IVT. The size and integrity of the biotinylated RNA were checked on a 6% Urea gel (Thermo Scientific, Darmstadt, Germany). Immediately before use, the RNA was heated to 95°C for 2min, on ice for 3min, and left at RT for 25min in RNA folding buffer (20mM Tris-HCl pH 7.5, 100mM KCl, 10mM MgCl2, 20U RNase OUT) to ensure correct folding. For each pulldown reaction, 1.5M hAoSMCs p6/p7 were used. Cells were washed twice with PBS before collection. After another wash, cells were lysed in lysis buffer (50mM Tris-HCl pH8, 150mM NaCl, 0.5% (v/v) NP40, 1x Roche cOmplete protease inhibitor, 20U RNaseOUT) for 30 min followed by a full-speed 30 min centrifugation. The supernatant was diluted in dilution buffer (20mM Tris-HCl pH 7.4, 150mM NaCl, 2mM EDTA pH 8.0, 0.5% Triton-X, and freshly added RNaseOUT and 2x Roche cOmplete protease inhibitor) and a pre-clearing step with washed MyC1 streptavidin beads (Invitrogen, Darmstadt, Germany) was performed. Beads were removed and a 5% Input sample was taken for control purposes. 250ng of biotinylated IVT *NUDT6* (or *GFP*, as a control reaction) were incubated with pre-cleared cell lysate on an overhead shaker at 4°C overnight. On the next day, the streptavidin beads were added to the samples for 3hrs on an overhead shaker at 4°C. Bead-target complexes were washed 3x in dilution buffer on the shaker at 4°C followed by one wash with 200mM NaCl – dilution buffer. For LC-MS analysis, samples were washed briefly in M/S buffer (10mM Tris-HCl pH 7.4, 150mM NaCl). To control the efficiency of the pulldown, both q-RTPCR and silver staining was performed.

### Liquid chromatography – Mass Spectrometry

Mass spectrometry data, together with a detailed description of methods and data analysis, have been deposited to the ProteomeXchange Consortium via the PRIDE partner repository (60) with the dataset identifier PXD033922. (Only for Reviewers: Username; Password: AQy4yigX).

## Statistics

Quantitative data are shown as dot plots with columns representing the mean and with error bars representing the SEM. Repetitive data are shown as dots representing the mean and with error bars representing the SEM connected by lines. Continuous variables between two groups were done by Student’s t-test. More than two groups were analyzed by 1-way ANOVA or 2-way ANOVA, depending on the number of independent variables. For repeated measures over time, multiple t-tests were used. P < 0.05 was used to show statistical significance with probability values two-tailed or one-tailed as indicated. Statistical analyses and graphs were generated with GraphPad Prism 8 or newer.

## Study Approval

All patients provided written and informed consent according to the Declaration of Helsinki. The study was approved by the local ethics committee.

Male *ApoE*^-/-^ mice (Taconic Biosciences, Hudson, NY, USA) and C57BL/6J mice (The Jackson Laboratory, Bar Harbor, ME, USA) were maintained in the local animal facility under ethical approval of the local ethics committee (Swedish Board for Agriculture; Ethical permit no. N48/16). The mice had *ad libitum* access to water and a standard chow diet. The animal study using mini-pigs was approved by the Animal Ethical Committee of the Government of Upper Bavaria (Munich, Germany; protocol No. ROB-55.2-2532.Vet_02-18-53) and performed in accordance with respective guidelines (EU Directive 2010/62/EU and Animal Welfare Act (2018)).

## Contributions

HW, FF, HJ, and LM designed the research studies. HW, GW, ABu, EC, FF, ABä, and HJ performed the experiments. ZW, BBK, DJvB, and IW acquired data. GW, ZW, BBK, DJvB, IW, and RB analyzed data and helped with data interpretation. NS and HHE provided resources in the form of human specimens. HW, HJ, and LM wrote the original draft of the manuscript. All authors reviewed and edited the manuscript.

## Acknowledgement

The authors thank Marie-Luise Engl, Nadiya Glukha, Renate Hegenloh, and Changyan Sun for their technical assistance.

This study is supported by the Swedish Heart-Lung-Foundation (20180680), the Swedish Research Council (Vetenkapsrådet, 2019-01577), the European Research Council (ERC-StG NORVAS), a DZHK Junior Research Group (JRG_LM_MRI), the SFB1123 and TRR267 of the German Research Council (DFG), the National Institutes of Health (NIH; 1R011HL150359-01), the Bavarian State Ministry of Health and Care through the research project DigiMed Bayern (all to L. Maegdefessel).

