## Supplemental Material for "Targeting long non-coding RNA *NUDT6* enhances smooth muscle cell survival and limits vascular disease progression"

### Supplemental Material and Methods

#### Patient Specimen

The tissue derived from the Munich Vascular Biobank (1). Carotid specimens were harvested during carotid endarterectomy, AAA samples were collected during open aneurysm repair and stored in DMEM/F12 medium (Sigma Aldrich, Darmstadt, Germany) for further processing (see section "Cell Culture and Transfection" for further details) cell culture purposes. Specimen for stainings were fixed in 2% zinc-paraformaldehyde for 48h followed by paraffin embedding. For characterization based on American Heart Association (AHA) staging after Stary (2) as well as fibrous cap thickness after Redgrave (3) , 5µm sections were stained with hematoxylin and eosin (HE) as well as Elastica van Gieson's (EvG). For RNA analysis, fresh frozen specimens stored at -80°C were used.

#### Immunohistochemistry

For frozen sections, the following primary abcam antibodies (Cambridge, UK) were used: a-Smooth muscle actin (ab5694), FGF2 (ab8880), Caspase 3 (ab13847), Ki-67(ab16667). As secondary antibody, HRP Goat Anti-Rabbit IgG (Vector Laboratories, Peterborough, UK) has been used. AEC staining (Histofine Simple Stain, Nichirei Bioscience Inc., Tokyo, Japan) has been performed under surveillance. Counterstaining with Mayer's Hematoxylin (Vector Laboratories, Peterborough, UK) for 10 seconds has been performed.

For FFPE sections, samples were rehydrated, and antigen retrieval was performed in pH6 in citrate buffer. H2O2 blocking was performed prior to primary antibody incubation. For this, the Dako REAL Detection System Peroxidase/DAB+, Rabbit/Mouse Kit (Dako Denmark, Glostrup, Denmark). The following steps were performed after the manufacturer's instructions. Slides were dehydrated in an increasing ethanol row followed by xylene. For mounting, EUKITT (Sigma Aldrich, Darmstadt, Germany) was used before imaging was performed.

#### RNA isolation and qRT-PCR

Cells were harvested 24h after transfection (described above). Homogenization of the cells was achieved by adding 700µL QIAzol (Qiagen, Hilden, Germany) to the cells, scraping them off with a cell scraper (Sigma Aldrich) and pipetting up and down three times. Tissues were lysed in 1mL QIAzol with a tissue homogenizer (ProScientific, Oxford, MS, USA). RNA isolation was performed with miRNeasy MiniKit (Qiagen, Hilden, Germany) following the instructions given by the provider. RNA concentration was measured using a Nanodrop 2000 spectrophotometer (Thermo Fisher Scientific, Gothenburg, Sweden). cDNA was synthesized with the cDNA RT kit (Applied Biosystems, Foster City, USA) according to the manufacturer's protocol using 10ng of RNA. TaqMan qRT PCR assay was performed for *NUDT6* and *FGF2* (Thermo Fisher Scientific, Gothenburg, Sweden) using 300-400ng of cDNA. Housekeeping control ACTB (Thermo Fisher Scientific, Gothenburg, Sweden) was used to normalize the assays. ΔΔCt was calculated in excel to determine the fold change. For more information about the used primers, see the Appendix.

**Cell culture and transfection**

Primary mouse aortic smooth muscle cells (mAoSMC), grown in DMEM/F12 (Gibco, Thermo Scientific, Gothenburg, Sweden) supplemented with 10%FBS (Gibco, Thermo Scientific, Gothenburg, Sweden) and 1% Penicillin-Streptomycin (PeSt) or human aortic/carotid smooth muscle cells (hAoSMC; hCtSMC), grown in SMC Growth Medium (PELO Biotech, Martinsried, Germany) were passaged up to 5-6 and seeded at 100,000 cells/well (6-well plate) or 10-15,000 cells/well (48-well plate).

Patient-derived cells from aneurysmatic or atherosclerotic tissue samples were collected during open AAA repair or carotid endarterectomy (CEA) and stored short-term in DMEM (Gibco, Thermo Scientific, Darmstadt, Germany) at +4°C. Then, the adventitia and -if possible – the endothelium was removed and the tissue was incubated at 37°C, 5%$CO_2$ on collagen-coated petridishes in SMC Growth Medium. Cells were used until passage 7.

Pig fibroblasts were provided by Assoc. Professor Yonglun Luo from Aarhus University, Denmark and derived from Yucatan minipigs.

*NUDT6* gene silencing and overexpression in hCtSMC and hAoSMCs were performed using Silencer™ Select Negative Control No. 1 siRNA (Ambion Life Technologies Corp., Carlsbad, CA, USA), *NUDT6* siRNA (Ambion Life Technologies Corp., Carlsbad, CA, USA), pCMV6-Entry Tagged Cloning Vector (OriGene Technologies, Inc. Rockville, MD, USA) and NUDT6 Human Tagged ORF Clone plasmid (OriGene Technologies, Inc. Rockville, MD, USA) according to the manufacturer's standard protocols. Lipofectamine RNAiMAX (Life Technologies Corp., Carlsbad, CA, USA) and Lipofectamine 3000 (Life Technologies Corp., Carlsbad, CA, USA) were used as transfection reagent. To further verify the effects of NUDT6 protein on HctSMCs, NUDT6 peptides were transfected with Pierce Protein Transfection Kit (Thermo Scientific Pierce RIPA Buffer, Thermo Scientific, Waltham, MA, USA) according to company's protocol. Pierce β-Galactosidase Control and Pierce FITC-Antibody Control in the transfection kit were used as control peptides for the experiment.

To further characterize the mechanism underlying enhanced *NUDT6*, cells were treated with 0.5 nM Ang II (Sigma Aldrich, Darmstadt, Germany), 20 µg/ml oxLDL (kindly provided by Eva Ehrenborg Group, Karolinska Institutet, Solna, Sweden) in Optimem (Gibco, Thermo Scientific, Darmstadt, Germany) supplied with 10% FBS, only Optimem supplied with 10% FBS, or Optimem supplied with 2% FBS as starvation group. RNA was isolated (see below) after 24 hours of treatment and kept in -80°C.

**RNA Immunoprecipitation (RIP)**

The Magna RIP 17-700 Kit (Millipore, Burlington, MA, USA) was used together with a polyclonal CRP1 antibody (PA5-86703, Invitrogen, Thermo Scientific, Darmstadt) to perform the RNA Immunoprecipitation according to the manufacturer's instructions. Briefly, hAoSMCs were grown in T175 flasks (1 flask per reaction and per analysis), washed with PBS, and lysed in the provided RIP lysis buffer (Millipore, Burlington, MA, USA) according to the protocol. The lysate was spun down at 12,000 rpm, and the supernatant was divided into 20% input and two reactions, CRP1 and IgG control. The supernatant was applied to the respective antibody-pre-incubated magnetic beads (5µg antibody per reaction). The provided rabbit IgG antibody was used as a control (Millipore, Burlington, MA, USA). After overnight incubation at 4°C, the reaction was washed 6x at 4°C before elution. The eluate as well as beads, input, and first wash supernatant were subjected to RNA and protein analysis.

**Protein extraction and western blotting**

HctSMCs or hAoSMCs were washed with PBS and resuspended in protein isolation buffer. Protein was then extracted for 30 minutes at 4°C in rotating mixer and quantified by using Pierce™ BCA Protein Assay Kit (Thermo Scientific, Waltham, MA, USA). 10 µg protein from each lysate were separated by 12% Mini-PROTEAN® TGX Stain-Free™ Protein Gels (Bio-Rad Laboratories AB, Solna, Sweden) in SDS-PAGE buffer in Mini-PROTEAN® Tetra Vertical Electrophoresis Cell (Bio-Rad Laboratories AB, Solna, Sweden). Gel was then transferred to a Trans-Blot® Turbo™ Mini polyvinylidene difluoride (PVDF) Transfer membrane (Bio-Rad Laboratories AB, Solna, Sweden) by using Trans-Blot® Turbo™ Transfer System (Bio-Rad Laboratories AB, Solna, Sweden), followed by blocking the membrane with 5% skim milk (Bio-Rad Laboratories AB, Solna, Sweden) (hCtSMCs) or 5% BSA (Sigma Aldrich) in TBS-T. Immunodetection of FGF2 and β-actin on membrane was performed through conjugated secondary antibodies to primer antibodies (Supplemental Table

2). Membrane was finally incubated in AmerSham™ ECL™ Prime western blotting detection reagent (GE Healthcare UK Ltd., Buckinghamshire, UK) for 5 minutes and exposed by CCD camera.

**Porcine In vivo Study**

8 female and castrated male LDLR-/- Yucatan minipigs (Exemplar Genetics, Coralville, IA, USA) were acclimatized prior surgery under conventional hygienic standards and received water ad libitum and pelleted high fat diet (Altromin, Lage, Germany).

PPE procedure:

On day 0, ultrasound and AAA induction by PPE procedure was performed, on day 8 – after ultrasound measurement - the aorta was treated with an endovascular balloon. On day 28, another ultrasound followed by euthanasia and tissue collection was performed.

Prior to anaesthesia, the animal was sedated (15mg/kg ketamine, 2mg/kg azaperone, 0.1mg/kg atropine) intramuscularly. Then, an intravenous catheter was placed and the anaesthesia was induced with 4-8mg/kg propofol. The animal was intubated (7-8.5mm cuffed endotracheal tube) and anaesthesia was maintained by continuously infusing 2.5-7mg/kg/h Propofol. Before skin incision, Cefuroxim (750mg) was administered and further analgesia was maintained throughout the surgery (50mg/kg metamizole, 4mg/kg carprofen, 0.001-0.01mg/kg fentanyl).

All ultrasounds were performed using a GE logiq S7 system (GE, Frankfurt, Germany) by two independent examiners. For orientation, the renal arteries and trifurcation was identified. Then, the aortic diameter was measured using leading edge technique in transverse and longitudinal sections at the point of maximum diameter.

The PPE procedure was performed via the left lateral flank. The aorta was isolated from the left renal artery to the aortic trifurcation. Lumbar arteries were clipped. Prior to aortic clamping, 3000IU Heparin was given. Depending on the measured aortic diameter, a 9-11x20mm PTA balloon (Medtronic Admiral Xtreme) was inserted and inflated to 10atm for one minute to secure pre-dilation of the aorta. By inserting a blunt needle (5G) into the aorta, the elastase perfusion (10IU/ml, porcine pancreatic elastase, Sigma Aldrich, Germany) could be performed for 10min using a pressure syringe. Then, the aorta was flushed with saline followed by closure of the aorta with a 5-0 suture. Closure of muscle, fat and skin layers was done and the wound was finally coated with a spray on dressing (Aqueos UK) to prevent infection.

Balloon Treatment:

The *NUDT6* GapmeR was prepared in a 33mM dilution (Qiagen, Hilden, Germany), spray coated on a 10- 12x20mm PTA balloon (Medtronic Admiral Xtreme) and introduced via the femoral artery advancing to the aneurysm which was identified *via* angiogram. The balloon then was inflated to 10atm for 3min. To exclude aortic occlusion or dissection, an angiography was performed.

Analgesia, Blood Sampling, Euthanasia:

Towards the end of anaesthesia, Buprenorphine (0.005-0.01mg/kg) was administered. In case of PPE procedure, administration was continued for at least 2 days. After the surgery, carprofen (4mg/kg) was administered orally for at least 5 days.

Blood sampling was performed at day 0,8 (+/-2) and 28 (+/-3) under anaesthesia.

On day 28, animals were sacrificed after a final aortic ultrasound using pentobarbital (>50mg/kg) and 40ml of 1M KCl solution.

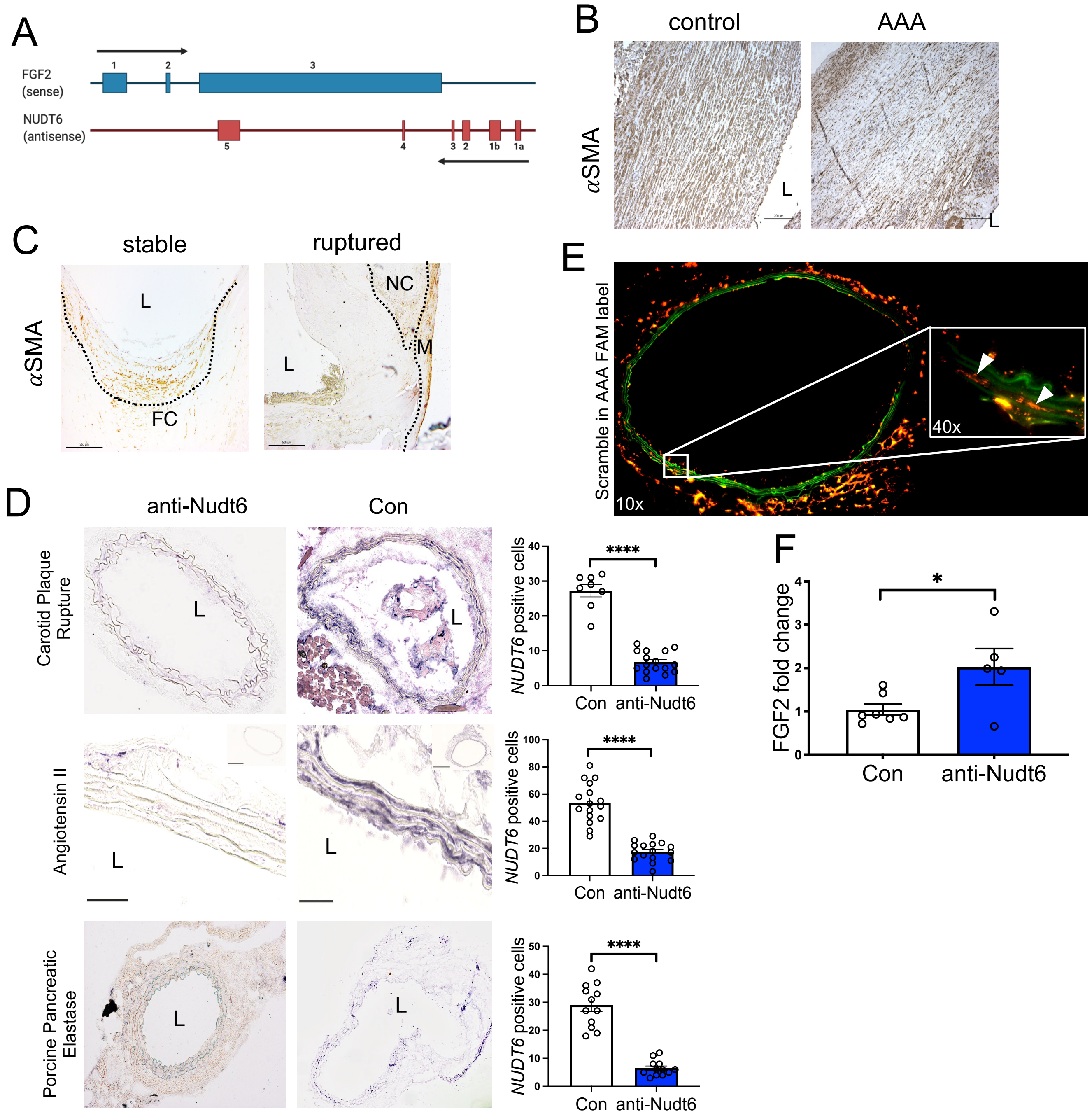


**Supplemental Figure 1**

**(A)** Scheme adapted from [24] showing the common overlap of *NUDT6* and *FGF2* transcripts and respective exons. **(B-C)** Immunohistochemical staining for $\alpha$SMA of stable and ruptured carotid lesions **(B)** and control aorta and AAA **(C)** show a similar expression pattern compared to FGF2. **(D)** Confirmatory *In situ* hybridization of Nudt6 in the three mouse models with signal quantification for *Nudt6* signal (4 high power fields per image, N=8-16 counts in total). **(E)** Fluorescent FAM-labelled scramble particles show successful transmission of locked nucleic acid – oligos to the intima media of the abdominal aorta *via* Ultrasound targeted microbubble destruction (UTMD). **(F)** qRT-PCR analysis of abdominal aorta show increased *Fgf2* levels in anti-*Nudt6* treated AngII mice (n=5 per group). Quantitative data are shown as mean + SEM. *p<0.05; ****p<0.0001. Significance is determined using one-tailed Student's t-test.

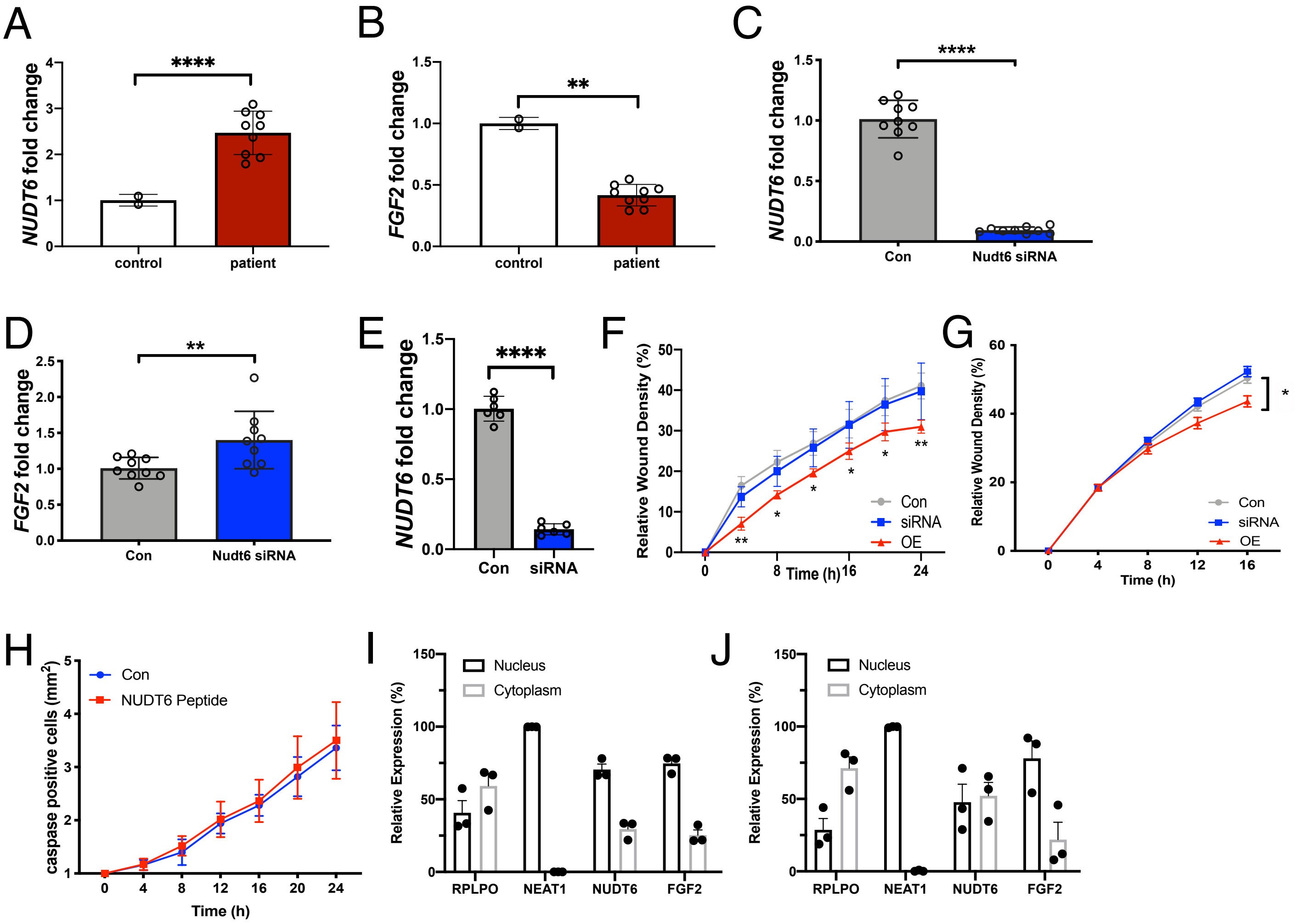


**Supplemental Figure 2**

**(A-B)** *NUDT6* **(A)** and *FGF2* **(B)** mRNA expression after qRT-PCR in 3 different patient-derived aortic VSMCs (n=3 per patient). **(C-D)** Knocking down *NUDT6* via siRNA in patient-derived aortic VSMCs led to a downregulation of *NUDT6* **(C)** mRNA but had a rescuing effect on *FGF2* **(D)** mRNA levels (n=3 per patient). **(E)** Confirmatory qRT-PCR for *NUDT6* after *NUDT6* knockdown shows significant downregulation in hAoSMCs (n=6 per group). **(F-G)** Live cell imaging of both hCtSMCs **(F)** and hAoSMCs **(G)** show impaired migratory capacity of cells receiving overexpression vector for *NUDT6*. *NUDT6-siRNA-treated* cells behave as scramble control (n=3 per group). **(H)** Dynamic live-cell imaging of hCtSMCs treated with NUDT6 peptide does not have an effect on apoptosis (n=5 per group). **(I-J)** Nucleocytoplasmic fractionation of hCtSMCs **(I)** and hAoSMCs **(J)** show intracellular distribution of *NUDT6* and *FGF2* compared to nuclear-expressed *NEAT1* and cytoplasmic expressed *RPLPO* (n=3 per group). Quantitative results are shown as mean+SEM. *p<0.05; **p<0.01; ****p<0.0001. Significance was calculated using One-tailed Student's t-test (**A-E, I-J**) or two-way ANOCA with Tukey **(F-H)**.

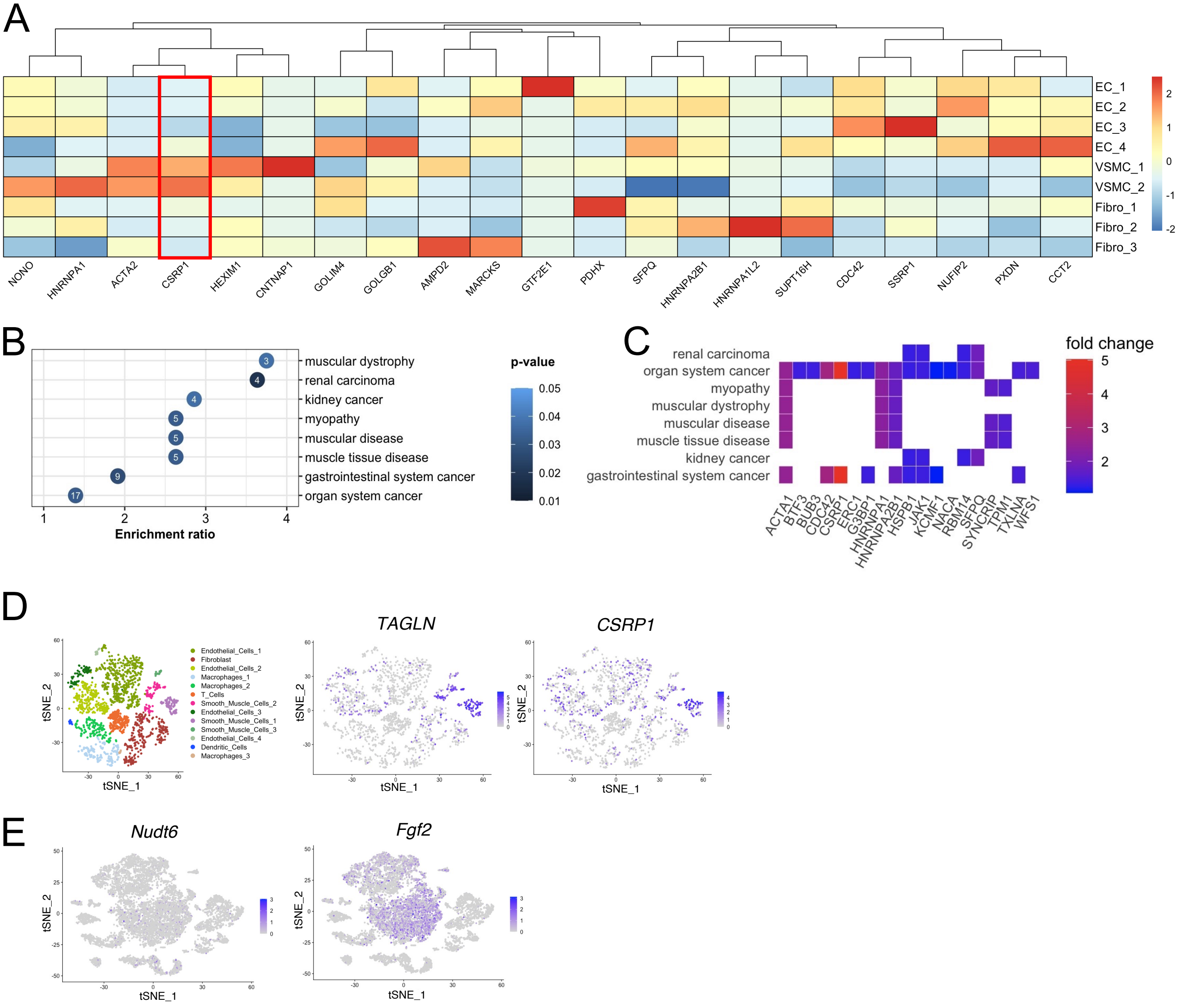


**Supplemental Figure 3**

**(A)** Heat map showing the abundance and expression of CSRP1 in VSMCs and ECs from human single-cell RNA sequencing experiments (Figure 5B). **(B)** Significant enrichment of potential NUDT6:protein interaction partners (Figure 5A) within several Disease Ontology datasets. **(C)** Regulation of potential NUDT6:protein interaction partners within specific Disease Ontology datasets shown in **(B). (D)** tSNE plots of scRNA Sequencing data from porcine AAA with expression plots of Transgelin (TAGLN) and CSRP1. AAAs were induced via PPE. **(E)** t-SNE plots of scRNA Sequencing data from murine AAA with expression plots of *Nudt6* and *Fgf2*.

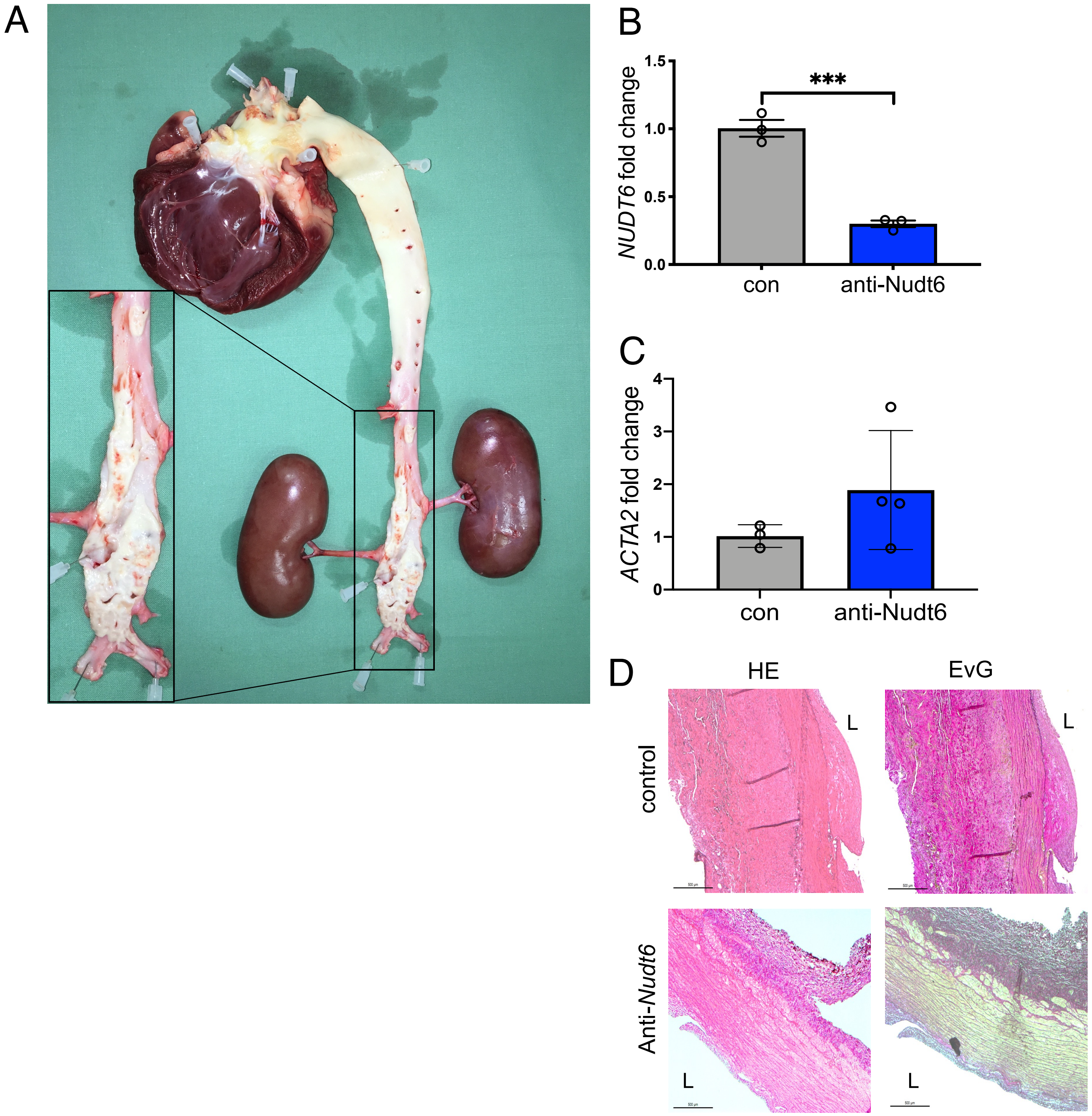


**Supplemental Figure 4**

**(A)** Representative image of the heart, aorta, and kidneys of an LDLR-/- pig with an enlarged image of the abdominal aorta. **(B)** Porcine aortic fibroblasts (n=3) treated with designed porcine *in vivo* locked nucleic acid against *NUDT6* show downregulation of *NUDT6* after treatment. © qRT-PCR of whole porcine aortic tissue (n=3-4) shows an increase in *ACTA2* mRNA. **(D)** HE and EvG stainings of the abdominal aorta anti-*NUDT6* treated (n=4) and control (n=3) animals. Quantitative data are shown as mean + SEM. ***p<0.001. Significance is determined using one-tailed Student's t-test **(B-C)**.

| | Protein names | Gene names | Fold enriched *NUDT6* vs CTRL |
|---|---|---|---|
| 1 | Cysteine and glycine-rich protein 1 | **CSRP1** | 32.588 |
| 2 | General transcription factor IIE subunit 1 | GTF2E1 | 11.126 |
| 3 | Cell division control protein 42 homolog | CDC42 | 6.958 |
| 4 | FACT complex subunit SPT16 | SUPT16H | 5.890 |
| 5 | Nuclear fragile X mental retardation-interacting protein 2 | NUFIP2 | 5.821 |
| 6 | Actin, alpha skeletal muscle;Actin, alpha cardiac muscle 1;Actin, gamma-enteric smooth muscle;Actin, aortic smooth muscle | ACTA1;ACTC1;ACTG2; ACTA2 | 5.646 |
| 7 | Peroxidasin homolog | PXDN | 5.446 |
| 8 | FACT complex subunit SSRP1 | SSRP1 | 5.404 |
| 9 | Heterogeneous nuclear ribonucleoprotein A1; | HNRNPA1;HNRNPA1L2 | 4.940 |
| 10 | T-complex protein 1 subunit beta | CCT2 | 4.879 |
| 11 | Protein HEXIM1 | HEXIM1 | 4.792 |
| 12 | Non-POU domain-containing octamer-binding protein | NONO | 4.729 |
| 13 | Myristoylated alanine-rich C-kinase substrate | MARCKS | 3.975 |
| 14 | Golgi integral membrane protein 4 | GOLIM4 | 3.930 |
| 15 | Contactin-associated protein 1 | CNTNAP1 | 3.741 |
| 16 | AMP deaminase 2 | AMPD2 | 3.662 |
| 17 | Splicing factor, proline- and glutamine-rich | SFPQ | 3.641 |
| 18 | Heterogeneous nuclear ribonucleoproteins A2/B1 | HNRNPA2B1;HNRPA2B1 | 3.421 |
| 19 | Golgin subfamily B member 1 | GOLGB1 | 3.384 |
| 20 | Pyruvate dehydrogenase protein X component, mitochondrial | PDHX | 3.331 |

**Supplemental Table 1**

Table of the top 20 identified *NUDT6*-RNA Pulldown Targets.
